# Ribosomal Protein bL27 Protects Translating Ribosomes from tmRNA-SmpB

**DOI:** 10.64898/2026.01.19.700400

**Authors:** Divyasorubini Seerpatham, George Wanes, Chathuri Pathirage, Mynthia Cabrera, Edu Usoro, Kristin Koutmou, Christine M. Dunham, Paul C. Whitford, Kenneth C. Keiler

## Abstract

Bacterial ribosomal protein bL27 is universally conserved and its amino terminus is adjacent to the peptidyl transfer center, yet its role in translation remains unclear. Combining genetics, biochemistry and molecular dynamics, we show that bL27 has an unexpected role in preventing *trans*-translation, the bacterial ribosome rescue mechanism, from interfering with protein synthesis. Deletion of the bL27 gene causes a 10,000-fold decrease in viability and this defect is partially rescued by deletion of the gene encoding tmRNA, a critical molecule for *trans*-translation. Molecular dynamics simulations also indicate that bL27 can slow the movement of tmRNA on the ribosome. These data link *trans*-translation and bL27, and support a model in which the amino terminus of bL27 acts as a gatekeeper to prevent tmRNA from sterically interfering with tRNA on the ribosome.

## Main Text

The genes encoding ribosomal protein bL27 (*rpmA*), tmRNA (*ssrA*), and SmpB (*smpB*) are among the set conserved in almost all bacterial species but absent from eukaryotes. tmRNA and SmpB function together in the *trans*-translation pathway to rescue ribosomes that are stalled at the 3’ end of an mRNA (non-stop ribosomes) (*1–3*). During *trans*-translation, the carboxyl terminal region of SmpB binds in the empty mRNA channel of a non-stop ribosome and the tmRNA-SmpB complex enters the decoding center of the ribosomal aminoacyl (A) site (*4–6*). A specialized reading frame within tmRNA is inserted into the mRNA channel, and translation resumes to add the tmRNA-encoded SsrA peptide to the end of the nascent polypeptide, ultimately releasing the ribosome at a normal stop codon (*7–12*). The rescue of non-stop ribosomes is essential in bacteria because damaged and truncated mRNAs are prevalent (*13*).

The single particle cryo-EM structure of a *trans*-translation inhibitor (KKL-2098) bound to a non-stop ribosome suggested that bL27 and tmRNA-SmpB may interact (*14–16*). KKL-2098 bound to a site near the peptidyl transfer center (PTC) that is not contacted by other antibiotics, and the bL27 amino terminal sequence adopted a novel position and was rotated ∼180° away from the PTC by bending at residue Gly8 (*14*). The amino terminal 8 residues of bL27 normally extend toward the PTC, making bL27 the protein closest to the catalytic site of the ribosome (*17*, *18*). Because of the proximity of bL27 to the PTC, bL27 was proposed to play a role in peptidyl transferase activity (*19–22*). However, more recent experiments show that removing bL27 from the ribosome does not change the catalytic rate (*23*). Mutation of the bL27 amino terminal sequence also has minimal effect on tRNA selection, the rate of elongation or the rate of peptide release during canonical translation (*23*). Nevertheless, deletion of bL27 in *Escherichia coli* (*E. coli*) results in slow growth. Since the protein is universally conserved in bacteria, this phenotype suggests that bL27 has a critical function in protein production (*24*, *25*). In addition, *E. coli* carrying amino terminal truncations of bL27 were hypersensitive to *trans*-translation inhibitors, indicating that bL27 and *trans*-translation may be linked (*14*, *15*). Here, we demonstrate that bL27 interferes with entry of tmRNA-SmpB into the PTC. Based on these data, we propose that a major role for bL27 is to prevent tmRNA-SmpB from interfering in normal protein synthesis.

### Mutations in bL27 specifically increase *trans*-translation and susceptibility to *trans*-translation inhibitors

Because the amino terminus of bL27 moves away from the PTC when KKL-2098 is bound, we hypothesized that this portion of bL27 might be required for *trans*-translation and inhibitors might function by displacing bL27 (*14*). To test these ideas, we mutated residues in the bL27 amino terminus or deleted bL27 entirely and measured antibiotic susceptibility and *trans*-translation activity. The mutant with no bL27 was hypersensitive to *trans*-translation inhibitors KKL-35 and MBX-4132 (which are acylaminooxadiazoles similar to KKL-2098; Fig. S1A), with IC_50_ values ∼50% of the values measured for strains with wild-type bL27 (Fig. S1B). Mutation of the broadly conserved HKK residues in bL27 (AHKKAGG to AaaaAGG; mutated residues shown in lower case) increased susceptibility to an extent comparable to removing bL27 entirely (Fig. S1B). In contrast, none of the bL27 mutants showed a statistically significant change in susceptibility to linezolid (Fig. S2C), which binds near the PTC, or to tetracycline (Fig. S2D), which binds the small subunit of the ribosome (*26–28*). The specific increase in susceptibility to acylaminooxadiazoles caused by bL27 mutations indicates that wild-type bL27 interferes with inhibitor efficacy.

To investigate the contribution of bL27 to *trans*-translation, we purified ribosomes from mutant strains and measured *trans*-translation in vitro with different concentrations of tmRNA-SmpB. Ribosomes from all strains produced SsrA-tagged DHFR, indicating that bL27 is not essential for *trans*-translation (Fig. 1A). Dose-response experiments showed that the concentration of tmRNA–SmpB required to achieve half-maximal tagging activity (EC_50_) on ribosomes with no bL27 or with the AaaaAGG mutant was <25% of the EC_50_ on ribosomes with wild-type bL27. These data indicate that tmRNA-SmpB interacted with the mutant ribosomes more efficiently than ribosomes with wild-type bL27 (Fig. 1B). Our results suggest that wild-type bL27 interferes with *trans*-translation and acylaminooxadiazole activity, in contrast to our initial hypothesis (*14*).

**Fig. 1.**
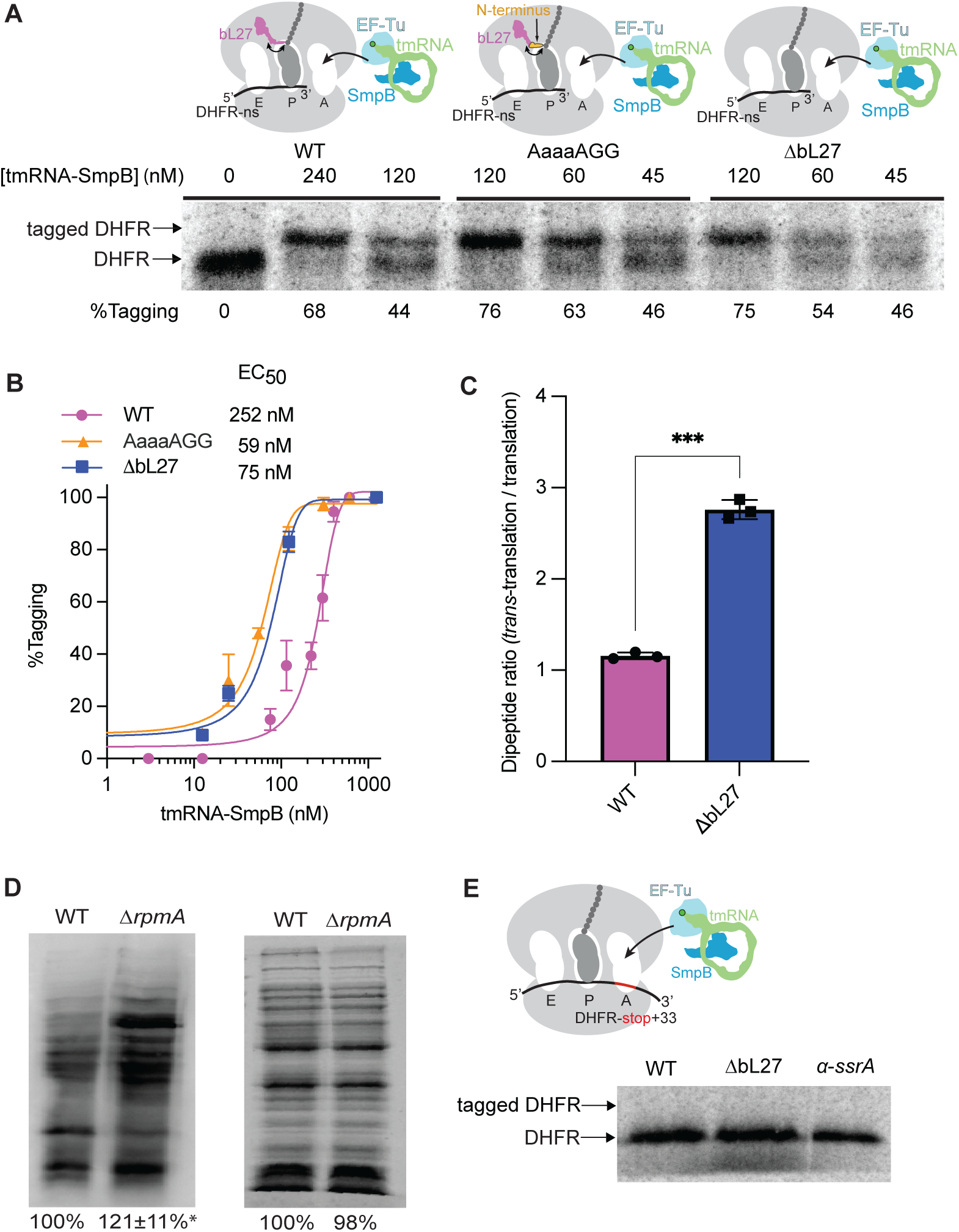
*trans*-Translation activity is higher on bL27 mutant ribosomes. In vitro *trans*-translation assays were performed using a non-stop DHFR template and wild-type, AaaaAGG, or ΔbL27 ribosomes, with varying concentrations of tmRNA-SmpB. **(A)** Gel image of a representative assay. The mobility of tagged and untagged DHFR is indicated with the percent tagging calculated after quantification of the tagged and untagged bands. **(B)** The average percent tagging from experiments as in (A) was plotted with whiskers indicating standard deviations. Averages were calculated from at least 3 trials at each concentration. The data were fit to a 3-parameter sigmoid equation and EC_50_ is shown. **(C)** Transfer of methionine from tRNA^fMet^ to tmRNA or tRNA^Ala^ in an in vitro system using wild-type or ΔbL27 ribosomes was monitored by eTLC and quantified. The *trans*-translation endpoint product was normalized to the translation endpoint product from the same ribosomes (*n*=3, \*\*\**P* value = 0.0005, Student’s unpaired *t*-test). **(D)** The amount of *trans*-translation in wild-type and Δ*rpmA* cells expressing tmRNA-HHHHHHD was measured by western blotting with an anti-his6 antibody. (Left) A representative western blot is shown. The total signal was quantified and normalized to wild-type. Averages from 3 repeats are indicated with the standard deviation (\**P* = 0.04 two tailed one-sample Student’s *t*-test vs 100%). (Right) Coomassie-stained gel showing the amounts of total protein loaded for the western blot analysis. **(E)** In vitro *trans*-translation assays were performed using DHFR-stop+33 template and no release factors with wild-type or ΔbL27 ribosomes. A representative gel is shown, with arrows indicating the mobility of the tagged and untagged DHFR. A control reaction containing an *α*-SsrA oligo to eliminate *trans*-translation is included to confirm the mobility of untagged DHFR.

To test if peptidyl transfer to tmRNA is altered in ΔbL27 ribosomes, we assembled wild-type and ΔbL27 ribosomes on non-stop mRNA with ^35^S-fMet-tRNA^fMet^ in the P site and added alanine-charged tmRNA-SmpB-EF-Tu-GTP complex to the A site (Fig. 1C, Fig. S2). The extent of peptidyl-transfer to tmRNA was compared to translation on a canonical, stop-codon containing transcript. We find that the similar levels of peptidyl-transfer are achieved for wild-type ribosomes undergoing “normal” and trans-translation. In contrast, ΔbL27 ribosomes exhibit a nearly 3-fold increase in peptidyl-transfer to tmRNA (Fig. 1C).

### bL27 is a gatekeeper for tmRNA-SmpB movement on the ribosome

The increased efficiency of *trans*-translation in vitro in the absence of bL27 suggests that the wild-type amino terminal sequence of bL27 may limit access of tmRNA-SmpB to the PTC. If that is true, there should be more *trans*-translation in cells lacking bL27. To test this, we expressed a tmRNA variant that encodes a peptide tag ending in HHHHHHD (*29*). This tag does not target proteins for degradation, so tagged proteins can be observed using an anti-His tag antibody. Western blots loaded with equal amounts of total protein from wild-type and Δr*pmA* strains showed that there are substantially more tagged proteins in the absence of bL27 (Fig. 1D).

Despite increased *trans*-translation efficiency on ΔbL27 ribosomes in vitro and in vivo, we find that *trans*-translation on these ribosomes still requires a truncated mRNA. We tested whether *trans*-translation could occur in the middle of an mRNA by repeating the in vitro assays using a DHFR template containing a stop codon at the 3’ end of the gene followed by a 33-nucleotide extension (DHFR-stop+33), while release factors were omitted from the reaction to prevent termination. Previous experiments showed that when a wild-type ribosome stalls at the stop codon on DHFR-stop+33 the mRNA extension occupies the mRNA channel, preventing *trans*-translation (*30*). Here, no *trans*-translation was observed with wild-type or ΔbL27 ribosomes on DHFR-stop+33 (Fig. 1E). These results suggest that substrate specificity is not radically altered in the ΔbL27 ribosomes and that the increased *trans*-translation observed in cells lacking bL27 is likely due to more efficient *trans*-translation on non-stop ribosomes.

Because tmRNA-SmpB interacts more efficiently with bL27 mutant ribosomes, we asked whether canonical translation is increased on mutant ribosomes by monitoring protein production over 90 min from a DHFR template that contained a stop codon. We found that ribosomes containing the AaaaAGG mutant bL27 produced approximately the same amount of DHFR as the wild-type ribosomes at all time points tested. This result indicates that mutation of the HKK residues of bL27 does not affect translation (Fig. 2A and B). However, with ribosomes that lacked bL27, the amount of protein synthesized was lower at 15 and 30 min, consistent with previous data (*23*). Previous studies with mutant strains of *E. coli* also found that deleting bL27 interfered with ribosome assembly (*1*). However, composite agarose-polyacrylamide gels and sucrose gradient profiles did not detect an increased fraction of partially assembled ribosomes in the *ΔrpmA* sample (Fig. 2C and D). Further, translation initiation assays indicated that the fraction of ribosomes that can form an initiation complex is the same in purifications of wild-type and *ΔrpmA* samples (Fig. 2E), suggesting that the decreased rate of protein synthesis in these assays is due primarily to slower translation in the absence of bL27 and not an increase in misassembled ribosomes.

**Fig. 2.**
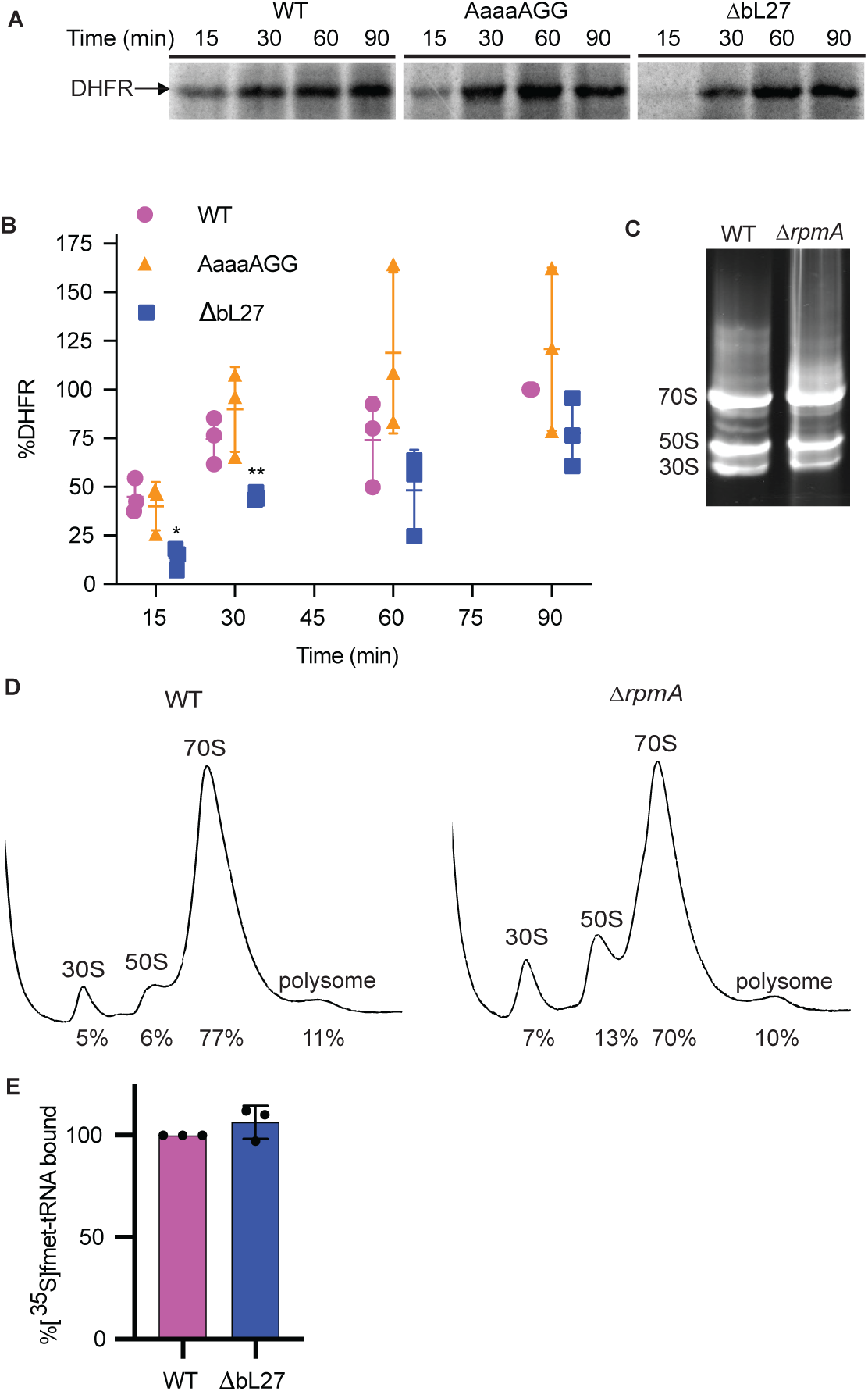
Effects of bL27 mutations on translation. **(A)** In vitro translation reactions were performed with wild-type and mutant ribosomes. A representative gel image is shown with an arrow indicating the mobility of the DHFR product. **(B)** Gels as in (A) were quantified, the amount of DHFR protein was normalized to the level produced by wild-type at 90 min and plotted with the mean and standard deviation indicated (*n* = 3: \**P* = 0.0064, \*\**P* = 0.0135 two-tailed Student’s *t*-test vs. WT). **(C)** Cell lysates from the indicated strains were separated on agarose–polyacrylamide composite gels to visualize ribosomal subunits. A representative gel is shown, with the mobility of 70S ribosomes and 50S and 30S subunits indicated. **(D)** Sucrose density gradient profiles on cell lysates from the indicated strains. Peaks were quantified and the percentage relative to total ribosomes is indicated. **(E)** The ability of purified ribosomes to form an initiation complex was measured by incubating ribosomes with DHFR mRNA, initiation factors, tRNA^fMet^ and [^35^S]Met, and quantifying radioactivity retained on nitrocellulose filters. The percentage of [^35^S]Met-tRNA bound to ΔbL27 ribosomes was calculated relative to wild-type and is not significantly different (*n* = 3).

We next used molecular simulations to gain insights into how bL27 can impact tmRNA movement through the ribosome. For these simulations, we used an all-atom structure-based (SMOG) model (*32*) to simulate movement of tmRNA-SmpB between the A and P sites. We specifically compare the dynamics the dynamics when the bL27 amino terminal tail is present or absent to determine the extent to which the steric composition of the tail can impact the kinetics of tmRNA movement.

These simulations indicate that the presence of the tail reduces accessibility of tmRNA to the P site, consistent with the in vitro and in vivo measurements. We simulated 150 independent events for each system and found that the dwell time of tRNA-like domain of tmRNA in the A site is increased by roughly 2-fold when the full-length bL27 protein is present (Fig. 3), which is comparable to the ∼4-fold difference in EC_50_ values (Fig. 1B). This delay may be understood as arising from steric interactions between the flexible amino terminus and the TLD acceptor stem (Fig 3C and D). That is, the amino terminal sequence bL27 can impede movement of the tmRNA, thereby delaying access to the P site (Fig. 3E and F).

**Fig. 3.**
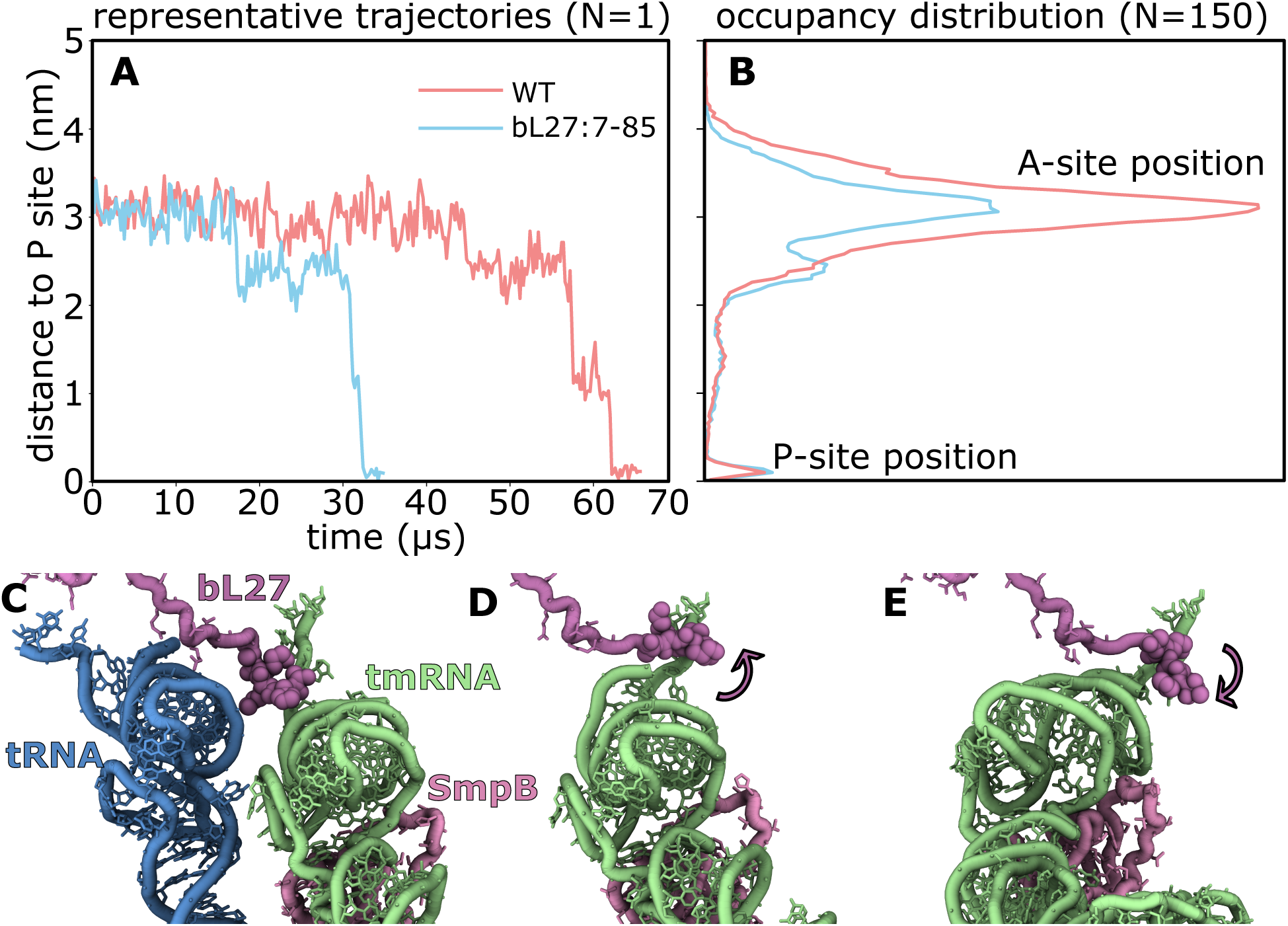
bL27 slows movement of tmRNA-SmpB during translocation. **(A)** Representative simulations of tmRNA-SmpB translocation with full-length (red) and truncated bL27 (blue) show that the N-terminal tail prolongs sampling of the ribosomal A site. The distance to P site is defined as the distance of the P atom of U9 from its post-translocation position on the LSU. **(B)** Occupancy of tmRNA position for the full-length and truncated systems, each calculated from 150 independent simulations. **(C)** Initially (distance ∼ 3 nm), deacylated tRNA (blue) leaves the P site and the disordered tail of bL27 (purple) makes close contact with tmRNA (green). **(D)** During tmRNA movement toward the P site (distance ∼2.2 nm), bL27 is sterically confined, which impedes the transition. **(E)** After tmRNA reaches the P site (distance ∼ 0.1 nm) the bL27 tail can again adopt an extended disordered form.

Taken together, these analyses suggest that the bL27 amino terminus can act as a dynamic steric obstacle that transiently restricts tmRNA motion between the A and P sites. This provides a physical explanation for the biochemical observation that *trans*-translation activity is lower with wild-type ribosomes.

### Deletion of *ssrA* rescues death phenotype in cells lacking bL27

Previous work with mutant *E. coli* strains has shown that bL27 is not essential but cells grow slowly without it (*24*, *25*). Using the wild-type strain MG1655 we found that deletion of bL27 leads to significantly slower growth due to two distinct effects (Fig. 4A). Plating assays showed that saturated cultures of Δ*rpmA* cells have a 10,000-fold decrease in colony forming units (CFU) compared to wild-type, indicating that most of the Δ*rpmA* cells are dead (Fig. 4B). This decrease in CFU causes an extended lag phase compared to wild-type. Once the Δ*rpmA* cells enter exponential growth they have a slower doubling time than wild-type. Both phenotypes were largely complemented by expression of bL27 from a multicopy plasmid (Fig. 4).

**Fig. 4.**
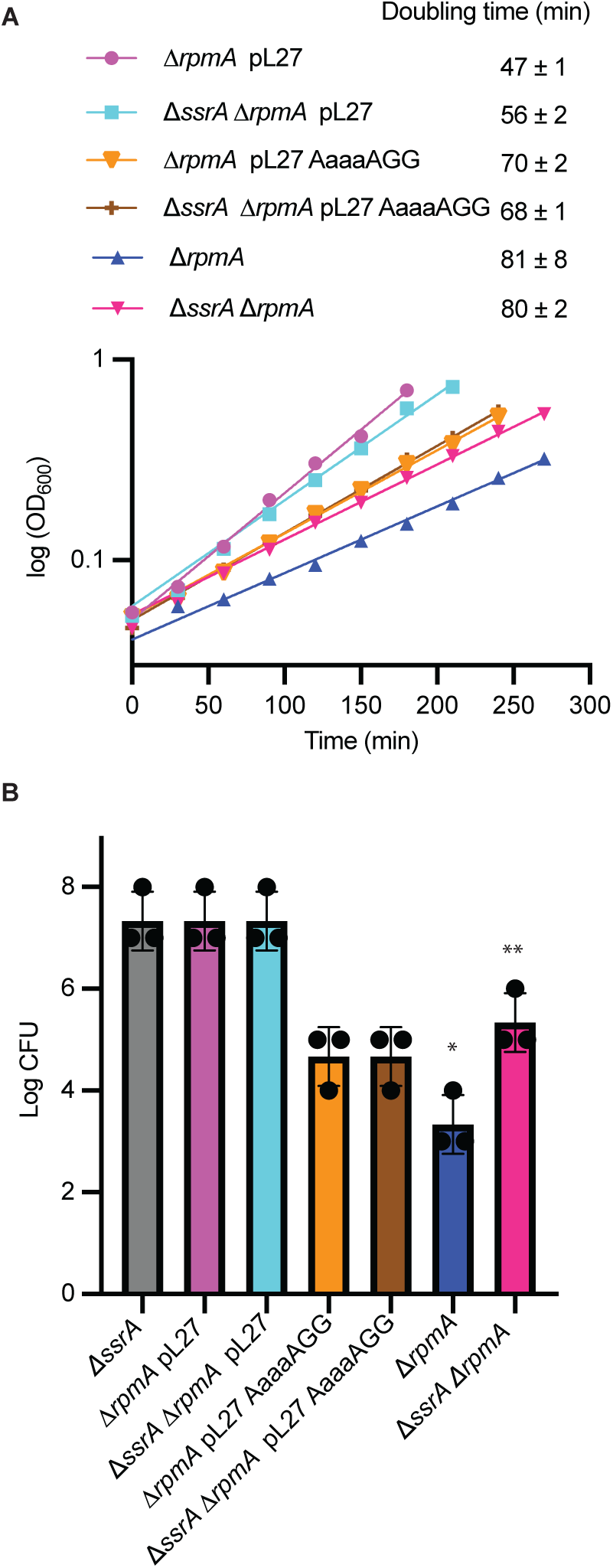
Deletion of *ssrA* partially complements the growth defect of Δ*rpmA* strain in stationary phase. **(A)** Growth curves of mutant strains monitored during exponential growth. The average doubling times (± standard deviation) from 3 experiments are shown. **(B)** Bar graph shows log CFU determined by spot plate method and deletion of *ssrA* in Δ*rpmA* strain had 100-fold higher CFU than Δ*rpmA* (*n* = 3 independent biological replicates; unpaired two-tailed Student’s *t*-test, \**P* = 0.25, \*\**P* = 0.26 compared to Δ*ssrA*).

To determine if *trans*-translation plays a role in the Δ*rpmA* growth defect, we deleted *ssrA* in the wild-type and Δ*rpmA* strains and measured growth parameters. Deletion of *ssrA* increases the CFU in Δ*rpmA* cells by 100-fold but has no detectable effect on the CFU for wild-type (Fig. 4B). In contrast, deletion of *ssrA* had little impact on the growth rate in exponential phase. These results indicate that tmRNA is problematic in the absence of bL27 when cells are in stationary phase, but not during exponential growth.

### bL27 limits tmRNA-SmpB interference with translation elongation

Why does tmRNA increase cell death in the absence of bL27? Our biochemical data can be explained by a model in which bL27 limits interaction of tmRNA-SmpB with elongating ribosomes as well as nonstop ribosomes. In this scenario, the absence of bL27 could allow tmRNA-SmpB to interfere with the dynamics of translation elongation. To test this possibility, we performed in vitro translation assays in the presence and absence of tmRNA-SmpB. As expected, addition of tmRNA-SmpB to reactions with wild-type ribosomes had little effect (Fig. 5A, B). However, with ΔbL27 ribosomes, addition of tmRNA-SmpB decreased the rate of protein production by 23% (Fig.5A, B). This result confirms that bL27 limits interference by tmRNA-SmpB with translation.

**Fig. 5.**
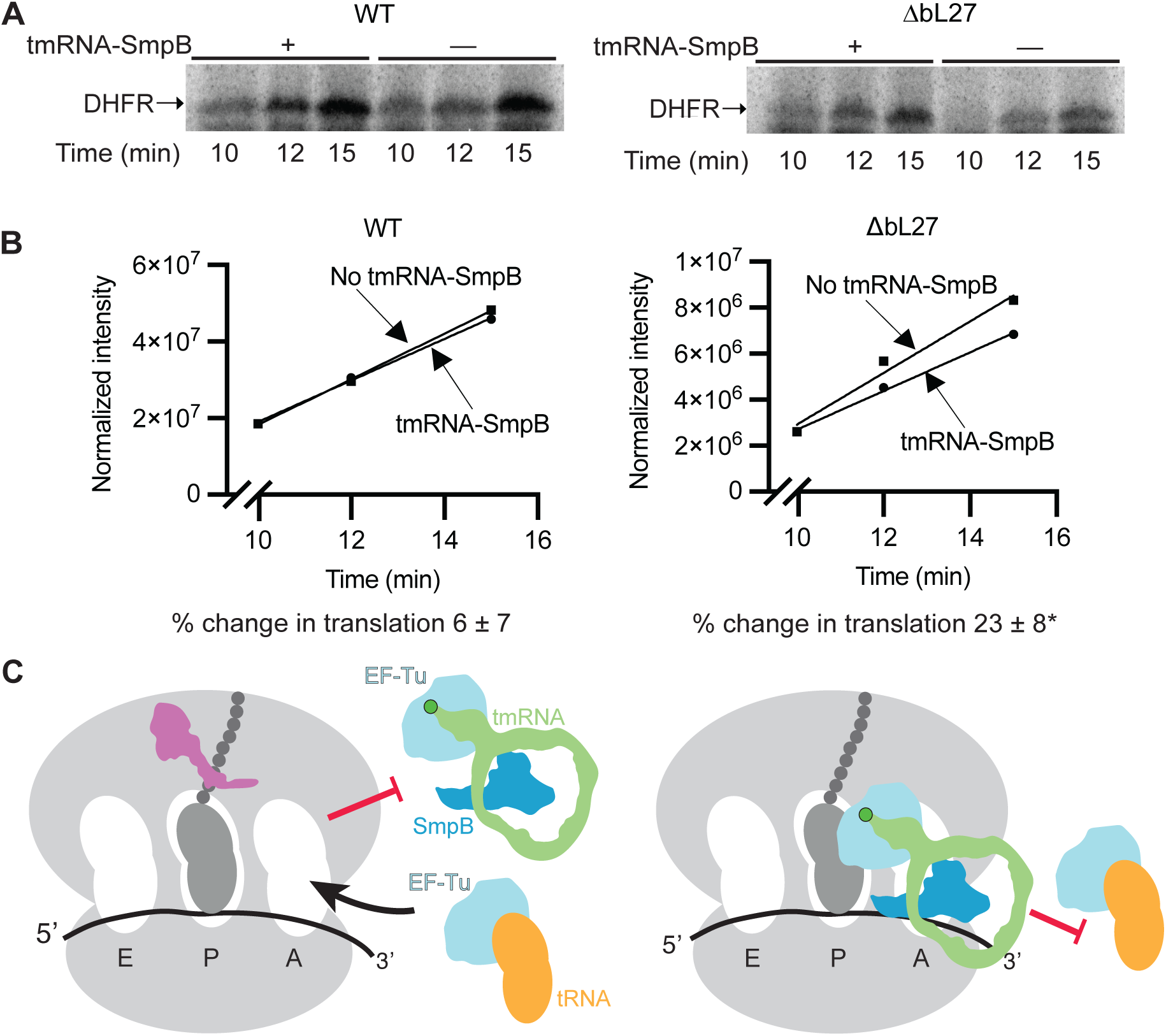
The rate of protein production decreases in the absence of bL27 with tmRNA-SmpB. **(A)** In vitro translation reactions were performed using the DHFR-Stop+33 template in the presence or absence of tmRNA-SmpB. Representative gels are shown. **(B)** Intensity of DHFR formed in the reaction vs time is plotted (representative curve). The initial rate of translation was calculated based on the slope of the curves, and the average percent change in translation rates with tmRNA-SmpB ± standard deviation of 3 trials is shown. (\**P* = 0.012 compared with wild type (two-tailed Student’s *t*-test)). **(C)** Model for the role of tmRNA-SmpB during elongation. bL27 limits access of tmRNA-SmpB to the PTC, allowing rapid entry of tRNAs to the A site. In the absence of bL27, tmRNA-SmpB has a longer dwell time in the A site and limits entry of tRNAs.

Our biochemical and computational analyses provide two possible explanations for how tmRNA-SmpB association with the ribosome leads to this reduced rate of protein production. First, during elongation tmRNA-SmpB may transiently interact with the A site of the ribosome, accommodating when there is no mRNA and dissociating when mRNA is present. In the absence of bL27 these transient events may be prolonged, which would reduce the rate of translation by sterically blocking tRNAs and RFs. Alternatively, the absence of bL27 may allow tmRNA to more easily access the PTC during elongation, which could allow for premature transpeptidation events, leading to inactive ribosomes. In either case, the data indicate a central role for bL27 in limiting interference by tmRNA-SmpB with translation elongation (Fig. 5C).

Both bL27 and tmRNA-SmpB are conserved in the vast majority of bacterial species, and the N-terminal residues of bL27 are almost invariant (Fig. S3). The functional interaction we observe here may account for this co-conservation. The absence of bL27 and tmRNA-SmpB from eukaryotes provides the opportunity to target this interaction for new broad-spectrum antibiotics.

## Acknowledgments

We thank Dan Herschlag, Bob Sauer, and Dan Kahne for insightful comments.

## Funding

National Institutes of Health grant R01GM12650 (KCK)

National Institutes of Health grant R35GM153502 (PCW)

National Science Foundation grant PHY2019745 (PCW)

National Institutes of Health grant R35GM128836 (KSK)

National Institutes of Health grant R35GM156629 (CMD)

We thank AMD for the donation of critical hardware and support resources from its HPC Fund that made this work possible.

We also acknowledge generous support from the Northeastern University Explorer cluster and the Northeastern University Research Computing staff.

## Author contributions

Conceptualization: KCK, PCW, KSK, CMD, DS

Investigation: DS, GW, MC, KCK, PCW CP, EU

Funding acquisition: KCK, CMD, PCW, KSK

Writing – original draft: DS, MC, KCK, GW, PCW

Writing – review & editing: DS, MC, KCK, GW, PCW, CMD, CP, KSK

## Competing interests

Authors declare that they have no competing interests.

## Data and materials availability

All data are available in the main text or the supplementary materials.

## Supplementary Materials

Materials and Methods

Supplementary Text

Figs. S1 to S3

Tables S1 to S2

## Notes

### Competing Interest Statement

The authors have declared no competing interest.

