## Supplemental materials for "Ribosomal Protein bL27 Protects Translating Ribosomes from tmRNA-SmpB"

**The PDF file includes:**

Materials and Methods
Supplementary Text
Figs. S1 to S3
Tables S1 to S2

### Materials and Methods

Bacterial strains, plasmids, and growth conditions

All strains (Table S1) were grown at 37 °C with shaking (250 rpm) in lysogeny broth (LB) supplemented with 50 μg/mL ampicillin, 30 μg/mL kanamycin, 20 μg/mL chloramphenicol, or 1 mM isopropyl-thio-*β*-d-galactoside (IPTG) as appropriate unless otherwise described.

To make the strain X90 *ssrA*6HisD Amp$^R$, Δ*rpmA*::kan, Δ*rpmA*::kan was moved from IW312 Δ*rpmA*::kan (Table S1) into X90 *ssrA*6HisD Amp$^R$ by P1 phage-mediated transduction. To delete Δ*rpmA* in MG1655 Δ*tolC*::cat, Δ*rpmA*::kan was moved from IW312 Δ*rpmA*::kan (Table S1) into MG1655 Δ*tolC*::cat (Table S1) by P1 phage-mediated transduction. To delete *ssrA* in the Δ*rpmA* background, first the CAT cassette was flipped out of MG1655 Δ*tolC*::cat Δ*rpmA*::kan with pCP20 (Table S1), selected on plain LB plates and grown at 40 °C resulting in MG1655 Δ*tolC* Δ*rpmA*::kan. The Δ*ssrA*::cat deletion from MG1655 Δ*ssrA*::cat was moved into MG1655 Δ*tolC* Δ*rpmA*::kan by phage mediated transduction resulting in MG1655 Δ*tolC* Δ*rpmA*::kan Δ*ssrA*::cat. *rpmA* mutations were introduced into linearized pL27 plasmid via Hifi Assembly (NEB, Ipswich, MA) using a 60 bp oligonucleotide (IDT, Coralville, IA) containing the mutations of interest (Table S2) to make the 11 plasmids containing *rpmA* mutations (Table S1). The 12 plasmids containing *rpmA* mutations including the one with wild-type *rpmA* (Table S1) were then transformed separately into MG1655 Δ*tolC*::cat Δ*rpmA*::kan resulting in strains expressing mutant bL27 in *rpmA* deletion background (Table S1). Plasmids pL27, pL27 AaaaAGG or vector only were transformed into *ssrA* deletion background strain (MG1655 Δ*tolC* Δ*rpmA*::kan Δ*ssrA*::cat) to express bL27 mutations in *ssrA* deletion background (Table S1).

Growth rate measurements

The doubling time of all the strains were measured by $OD_{600}$ readings of each strain grown in 2 mL LB with required supplements. Starting $OD_{600}$ was set to 0.05 by inoculating from overnight cultures. Doubling times were calculated from the exponential phase ($OD_{600}$ = 0.1–0.6) fitting the curves to exponential growth curve function in GraphPad Prism (Version 10.6, GraphPad Software, LLC, San Diego, CA).

Spot plate assays

Overnight cultures were back diluted to $OD_{600}$ = 0.05 and grown for 48 h. Cultures were serially diluted tenfold in sterile LB medium. Aliquots (5 μL) of each dilution were spotted onto LB agar plates with the supplements, as needed. Plates were incubated at 37 °C for 24 h.

Growth inhibition measurements

Overnight cultures were diluted to $OD_{600}$ = 0.004 and added to 2-fold serial dilution concentrations of compounds or drugs in a 96-well plate, grown at 37 °C for 24 h, and $OD_{600}$ measured using a Spectramax plate reader (Molecular Devices, Sunnyvale,CA). The compound concentrations were plotted against normalized $OD_{600}$ in Sigmaplot v.10, San Jose, CA and fit with 3 parameter-sigmoidal function to obtain $IC_{50}$ values. Averages of $IC_{50}$ values with standard deviations and number of trials are indicated in Fig. S1. Statistical significance was tested using GraphPad Bonferroni Multiple Comparison ANOVA on selected pairs.

Purification of ribosomes for in vitro assays and sucrose density gradient profiles on lysates

Crude ribosomes were prepared as described previously *(33)*. 70S ribosomes were isolated by sucrose density fractionation. Crude ribosome (200 µl of 50 $A_{260}$ units) was loaded on a 10-40% linear gradient of sucrose in ribosome resuspension buffer (20 mM HEPES-KOH [pH 7.6], 60 mM $NH_4Cl$, 12 mM $MgCl_2$, 0.5 mM EDTA, 6 mM *β*-ME) and centrifuged at 151,000 *g* for 2.5 h in a Beckman SW-41 rotor (Beckman Coulter, Brea, CA). Fractions containing 70S ribosomes were harvested by fractionation with Piston Gradient Fractionator (Biocomp, Tatamagouche, NS, Canada) and fractions were pooled and centrifuged for 24 h at 220,000 *g* at 4 °C. The 70S ribosomal pellet was resuspended and stored at -80 °C in ribosome resuspension buffer. For sucrose density gradient profiles on lysates, cleared cell lysate (200 µl of 50 $A_{260}$ units) was loaded on a 10-40% sucrose linear gradient and fractionation was done as described above.

Isolation of tmRNA and purification of SmpB

*E. coli* tmRNA was transcribed in vitro as described previously (*34*). For purification of SmpB, *E. coli* BL21 (DE3) with pET28 *smpB* (Table S1) were grown in terrific broth supplemented with 50 µg/mL kanamycin at 37 °C to $OD_{600}$ = 0.6 and induced with 1 mM IPTG for 3 h. The cells were harvested by centrifugation at 6953 *g* for 20 min, resuspended in lysis buffer/wash buffer (100 mM $Na_2H_2PO4$ [pH 7.6], 150 mM KCl, 6 M GuHCl, 10 mM imidazole) lysed by sonication, and cell debris was removed by centrifugation at 28,000 *g* for 20 min. Cleared lysate was incubated with HisPur Ni-NTA agarose resin (Thermo Fisher Scientific, Waltham, MA,) for 1 h at 4 °C, and washed 4 times with 40 mL wash buffer, and bound protein was eluted with 10 mL of elution buffer (100 mM $Na_2H_2PO_4$ [pH 7.6], 300 mM KCl, 6 M GuHCl and 1 M imidazole). The eluate was dialyzed against buffer (50 mM HEPES-KOH [pH 7.6], 300 mM KCl, 10 mM $MgCl_2$, 20% (v/v) glycerol, and 7 mM *β*-ME) at 4 °C.

In vitro *trans*-translation and translation

*E. coli* in vitro transcription-translation system was designed based on the OnePot PURE system (*35*). The energy solution was prepared as described previously (*35*). tRNAs for energy solution was purified from *E. coli* MRE 600 as described previously (*36*). Protein solution was made with proteins purified as the same way as described previously but from strains listed in (Table S1) and was made without adding release factors (RF1, RF2, RF3) and EF-Tu (*35*). Release factor mix containing RF1, RF2 and RF3 was purified using strains listed in Table S1 as described previously (*35*). EF-Tu was purified as described previously (*37*). DNA template encoding dihydrofolate reductase (DHFR) gene without a stop codon (DHFR-ns), DNA template DHFR with a stop codon (DHFR-stop) and DNA template with DHFR with stop codon and 33 nucleotides past the stop codon (DHFR-stop+33) were prepared via PCR, as described previously (*34*, *38*). In vitro *trans*-translation reactions were set up with DHFR-ns template. The *trans*-translation reactions contained energy solution (2 µL), protein solution (1 µL), EF-Tu (10 µM), 70S ribosomes (2.2 µM) (wild-type ribosomes, AaaaAGG mutant bL27 containing ribosomes or ΔbL27 ribosomes), DHFR-ns DNA template (9 ng/µL), release factor mix (2.2 µM), [$^{35}$S]-methionine (0.42 µCi/µL) and varying equimolar concentrations of tmRNA and SmpB. Control reaction was set up the same way but with no tmRNA and smpB and to inhibit any background *trans*-translation activity 0.5 µM *α-ssrA* oligonucleotide was added (Table S2). The reactions were incubated at 37 ºC for 90 mins, acetone precipitated, boiled at 95 °C for 10 mins with 2× sample loading buffer (2% SDS, 20% glycerol, 0.004% bromophenol blue, 0.125M Tris-Cl, pH 6.8, 6 mM *β*-ME) analyzed by SDS-PAGE, and visualized by phosphor imaging (GE

Healthcare, Chicago, IL). The band intensity of tagged DHFR-ns (higher molecular weight due to addition of the SsrA tag) and DHFR-ns untagged were quantified using ImageJ. %*trans*-translation was calculated using the equation [tagged DHFR-ns / (DHFR-ns tagged + DFHR-ns untagged)]. [tmRNA-SmpB] added to the *trans*-translation reactions were plotted against %tagging and $EC_{50}$ was calculated in Sigmaplot v.10.

In vitro translation experiments were set up with the following modifications to the *trans*-translation assay. Translation was tested on DHFR-stop DNA template. 4X reaction were set up and incubated at 37 °C and samples was taken out at 15, 30, 60 and 90 mins and acetone precipitated and analyzed on SDS PAGE as described above. The band intensity of DHFR-stop was quantified. Relative %DHFR formed at each time point was calculated using the equation (DHFR-stop intensity / DHFR-stop intensity with wild-type ribosomes at 90 min).

To assess the competition of tmRNA and tRNA with or without bL27, in vitro translation reactions were set up with DHFR-stop+33 DNA template as described above with following modifications. The translation reactions were set up with and without tmRNA-SmpB (2.2 µM) and incubated at 30 °C and samples were withdrawn at 10, 12 and 15 mins for analysis. The intensity of DHFR band was quantified and plotted against time and the rate of translation was calculated from the slope of the plots. The percentage change in rate of translation was calculated as a percentage relative to rate of translation without tmRNA-SmpB.

#### Reconstituted in vitro translation assay

A reconstituted in vitro translation assay was used to measure the amount of $^{f}$Met-Ala dipeptide formed with tRNA or tmRNA in ribosome complexes with and without bL27. The ribosomes were purified as mentioned above. *E. coli* initiation factors, elongation factors, total tRNA and tRNA$^{fMet}$ were purified as previously described (*39*). 70S initiation complexes with $^{35}$S-$^{f}$Met-tRNA$^{fMet}$ positioned in the P site were assembled on either Z4C_A_GCC mRNA (sequence: GGCAAGGAGGUAAAAAUGGCC) for tRNA$^{Ala}$ or ns-mRNA (sequence: GGCAAGGAGGUAAAAAUG) for tmRNA. The mRNAs were purchased from IDT. Equal volumes of ternary complex containing EF-Tu, 5 mM GTP and either 6 µM aminoacylated tRNA$^{Ala}$ or 6 µM tmRNA and equimolar SmpB were added to pelleted initiation complexes (60 nM). tRNA and tmRNA were aminoacylated by *E. coli* Ala-tRNA aminoacyl synthetase (AlaRS). Reactions were quenched at 300 s by 500 mM KOH (final concentration). $^{35}$S-$^{f}$Met reactants were separated from $^{35}$S-$^{f}$Met-Ala products by electrophoretic TLC and visualized by phosphor imaging. Images were quantified with ImageQuant (GE, Boston, MA).

#### In vivo *trans*-translation tagging with or without bL27

To assess total in vivo tagging in strains with or without bL27, X90 *ssrA*6HisD and X90 *ssrA*6HisD Δ*rpmA*::kan (Table S1) were grown to an $OD_{600}$ of 0.6, harvested by centrifugation, and weighed. Pellets were resuspended in 2× sample loading buffer in proportion to wet cell mass and heated at 95 °C for 5 min. Western blotting was performed with Penta-His antibody (Qiagen, Hilden, Germany, 1:10,000 dilution) followed by goat anti-mouse IgG–alkaline phosphatase conjugate (Sigma-Aldrich, St. Louis, MO, 1:20,000 dilution). Signals were detected using ECF substrate (Cytiva, Amersham, Marlborough, MA) and visualized with ChemiDoc imaging system (Bio-Rad, Hercules, CA).

#### Translation initiation assay

Energy solution for translation initiation reaction was prepared as described previously with modifications (*35*). The energy solution included only methionine instead of all 20 amino acids and tRNAs were not added. A modified protein solution was prepared by purifying proteins as described previously but the modified protein solution contained proteins that are only required for translation initiation (creatine kinase, myokinase, nucleoside–diphosphate kinase, methionyl-tRNA transformylase, IF1, IF2 and IF3). DHFR-stop mRNA was in vitro transcribed as described previously (*35*). Met tRNA was in vitro transcribed as described previously with metZ DNA template with T7 promoter (*34*). metZ DNA template for in vitro transcription was amplified from *E. coli* genomic DNA using primer pairs listed in Table S2. In vitro translation initiation reactions were set up with 1 μL of modified energy solution, 0.8 μL of modified protein solution, 70S ribosomes (2 μM), DHFR-stop mRNA (2.5 μM), [35S]-methionine (0.42 μCi/μL) and $tRNA^{met}$ (2.5 μM). The reactions were incubated for 15 mins at 37 $^{\circ}$C. The reaction mix was passed under vacuum through 0.45 μm nitrocellulose membrane filter (MilliporeSigma, Burlington, MA) presoaked with wash buffer (20 mM HEPES-KOH [pH 7.6], 60 mM $NH_4Cl$, 12 mM $MgCl_2$, 50μg/ml Bovine Albumin Serum). The membranes were washed and dried, and radioactivity retained on the membrane was determined by scintillation counting.

Agarose–polyacrylamide composite gel electrophoresis

Cells were grown to $OD_{600}$ = 0.6 and lysed by sonication with buffers described previously (*33*). The lysate was cleared by centrifugation at 28,000 *g* for 30 min at 4 °C. Ribosome complexes were analyzed using agarose–polyacrylamide composite gels prepared in TBM buffer. Gels containing 1% agarose and 6% acrylamide (19:1) were pre-run for 60 min at 100 V in 1× TBM at 4 °C. Cleared lysates (10 $A_{260}$ units) were mixed with loading buffer (20% sucrose, 1× TBM, bromophenol blue) and resolved at 100 V for 180 min at 4 °C. Gels were stained with ethidium bromide (25 μg/mL) and visualized with UV illumination.

Molecular dynamics simulations

To investigate the potential impact of steric interactions between bL27 and tRNA-like domain (TLD) of tmRNA, we used a structure-based force field ("SMOG") of the tmRNA–SmpB–ribosome complex (*40*). In this approach, all tmRNA-SmpB interactions with the ribosome that are present in the post-translocation configuration (PDB 7ACJ, (*41*)) were explicitly defined as stabilizing, while intra-ribosomal contacts were defined to stabilize the pre-translocation conformation (PDB 7AC7, (*41*)). Non-native interactions were assigned repulsive potentials, which ensures that only sterically accessible conformations are sampled in the simulations. Since the stabilizing interactions are defined based on the endpoints (i.e. tmRNA in the A and P sites), this force field is suited to capture steric and entropic contributions to the dynamics (*32*, *42*, *43*). For example, these models have shown how tRNA dynamics is regulated by steric factors during P/E hybrid formation and translocation of tRNA (*44–46*). Here, to isolate the contributions of the bL27 N-terminal sequence, two sets of simulations were performed. In the first set, the bL27 N-terminal residues (Ala2-Ala6) were included and treated as disordered, consistent with the frequent lack of density for this region (PDB ID: 7ACJ). In the second set of simulations, we used an identical model, except that the N-terminal residues were removed (bL27:7-85).

Potential energy function

An all-atom structure-based (SMOG) force field of the tmRNA–SmpB–ribosome complex was employed. The potential energy may be expressed as:

$$\boldsymbol{U} = \sum_{\text{bonds}} \frac{\epsilon_r}{2}\left(r_i - r_{i,0}\right)^2 + \sum_{\text{angles}} \frac{\epsilon_\theta}{2}\left(\theta_i - \theta_{i,0}\right)^2 + \sum_{\text{impropers}} \frac{\epsilon_{\chi_{\text{imp}}}}{2}\left(\chi_i - \chi_{i,0}\right)^2$$
$$+ \sum_{\text{planars}} \frac{\epsilon_{\text{planar}}}{2} \cos(2\chi_i - \pi) + \sum_{\text{backbone dihedrals}} \epsilon_{\text{bb}} F\left(\phi_i - \phi_{i,0}\right)$$
$$+ \sum_{\text{sidechain dihedrals}} \epsilon_{\text{sc}} F\left(\phi_i - \phi_{i,0}\right) + \sum_{\text{intra-contacts}} \epsilon_c \left[\left(\frac{\sigma_{ij}}{r_{ij}}\right)^{12} - 2\left(\frac{\sigma_{ij}}{r_{ij}}\right)^{6}\right]$$
$$+ \sum_{\text{inter-contacts}} \frac{\epsilon_c}{3} \left[\left(\frac{\sigma_{ij}}{r_{ij}}\right)^{12} - 4\left(\frac{\sigma_{ij}}{r_{ij}}\right)^{3}\right] + \sum_{\text{non-contacts}} \left(\frac{C_{18}}{r_{ij}^{18}} - \frac{C_{12}}{r_{ij}^{12}}\right)$$

where

$$F(\phi) = [1 - \cos(\phi)] + \frac{1}{2}[1 - \cos(3\phi)]$$

$\{r_0\}$ and $\{\theta_0\}$ parameters are assigned according to the Amber99sb-ildn force field (*47*), whereas the dihedral parameters ($\{\chi_0\}$ and $\{\phi_0\}$) are given the corresponding values found in the experimental model. The energetic weights are $\epsilon_r = 100\ \frac{\epsilon}{\text{Å}^2}$, $\epsilon_\theta = 80\ \frac{\epsilon}{\text{rad}^2}$, $\epsilon_{\chi_{imp}} = 10\ \frac{\epsilon}{\text{rad}^2}$, $\epsilon_{\chi_{planar}} = 40\ \frac{\epsilon}{\text{rad}^2}$,where $\epsilon$ denotes the reduced energy unit. Non-bonded contacts found in the experimental model are identified using the Shadow Contact Map algorithm (*48*), with a shadowing radius of 1 Å and cutoff distance of 6 Å. Intramolecular contacts within the ribosome and the tmRNA–SmpB complex were modeled using an attractive 6–12 potential to stabilize a preassigned structure. The intra-ribosomal contacts and dihedrals were assigned to stabilize the pre-translocation structure (PDB ID: 7AC7) while the intra-tmRNA-SmpB complex contacts and dihedrals were defined based on the translocated structure (PDB ID: 7ACJ). To define the contacts between the ribosome and tmRNA–SmpB, long-range interactions were modeled with a 3–12 potential with the minimum assigned from the translocated configuration.

Since contact terms in structure-based model describe effective interactions (*49*), subsets of the contact weights were rescaled. Contacts between the head and body of 30S ribosomal subunit and between the head of the 30S and the large subunit were rescaled by a factor of 0.5 to allow for the head rotation and tilting (*50*) while interactions between A-site finger and 30S were rescaled by 0.2 to allow for the A-site finger to move towards its post-translocation position. Inter-complex contacts between tmRNA-SmpB and the ribosome were rescaled by a factor of 0.6 to account for the transient nature of tmRNA-SmpB binding to the ribosome.

Atom pairs not in native contact in the experimental structure were assigned a purely repulsive interaction to represent excluded-volume effects using the 12-18 potential. The weights of dihedral and contact energies were normalized following the procedure in as described previously (*51*).

In these simulations, we additionally introduced a harmonic potential (weight of $2\,\frac{\epsilon}{\text{Å}^2}$) between the 3'-CCA tail of the tmRNA and the P site of the ribosomal large subunit (LSU). Specifically, the restraint was introduced between the phosphate atoms of A363 in the tmRNA and C2064 in the LSU. This was introduced in order to mimic a post-peptidyltransfer state, where the CCA end is chemically bonded to the nascent protein chain. Accordingly, our model is suited to address the impacts of bL27 on translocation of tmRNA-SmpB, rather than probe the direct impact on transpeptidation.

Simulation details

All force field files were generated using the SMOG 2 (*40*) software package. Molecular dynamics simulations were performed using the OpenMM (*52*) and OpenSMOG (*53*) libraries. The system was simulated at a reduced temperature of $0.5\,\frac{\epsilon}{K_B}$ which was maintained using Langevin dynamics protocols with a timestep of 0.002. This temperature was chosen since it yields predictions of molecular flexibility that are consistent with explicit-solvent simulations and crystallographic B-factors for molecular assemblies (*17*). To estimate the effective simulated time, we defined one reduced time unit to correspond to approximately 1 ns, as estimated for tRNA motion within the ribosome (*55*). This estimate was based on diffusion coefficients calculated from simulations using both SMOG and explicit-solvent models.

In preliminary simulations with our model, it was observed that the TLD always reached the P site before the mRNA-like domain. In addition, as sterically required, the deacylated tRNA was always displaced from the P site prior to TLD movement to the P site. To probe the effects of bL27, we focused the current analysis on movement of the TLD.

Two sets of simulations were performed, each consisting of 150 independent runs: one with the complete bL27 N-terminal tail and the other with a truncated variant lacking residues Ala2–Ala6 (bL27:7-85). Each simulation was continued for up to $3.2 \times 10^8$ time steps. Simulations were continued until the TLD associated with the P site. For the full-length system, the TLD reached the P site in 112 simulations within the simulated time, while there were 130 successful events for the bL27:7-85 system. All 150 trajectories were used to calculate each occupancy distribution. Accordingly, the total occupancy values represent lower-bound estimates. However, since there was a larger number of simulations of the full-length system in which the tmRNA did not reach the P site, it is expected that the A-site occupancy will increase more for the full-length system, which would further amplify the differences described in the main text.

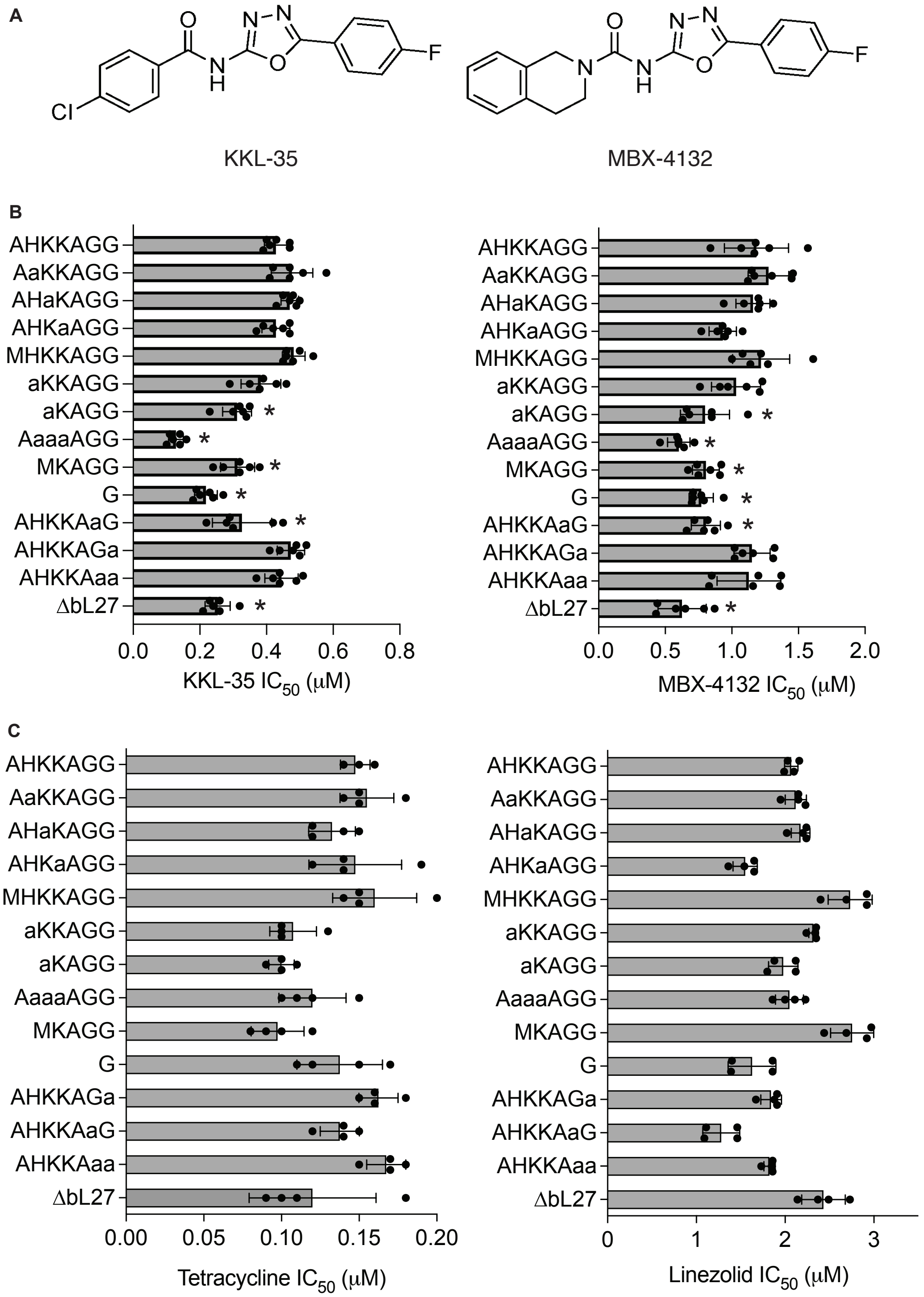


**Fig. S1. Antibacterial activity of small molecule inhibitors on bL27 strains. (A)** Structures of KKL-35 and MBX-4132. **(B)** The $IC_{50}$ for growth inhibition of strains with mutant bL27 by *trans*-translation inhibitors and other antibiotics were measured and the average is plotted with error bars indicating the standard deviation. **P* ≤ 0.03 compared with AHKKAGG (one-way ANOVA).

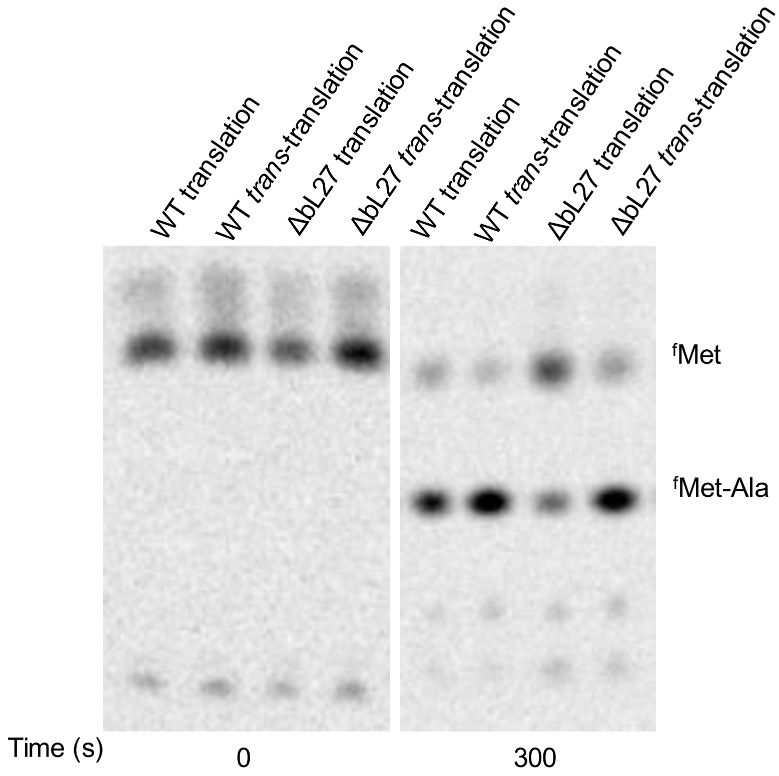


**Fig. S2 Representative eTLCs showing dipeptide formation in wild-type and Δbl27 ribosomes.** $^{f}$Met reactant at 0 s (left) and unreacted $^{f}$Met and $^{f}$Met-Ala product at 300 s (right) separated on eTLC.

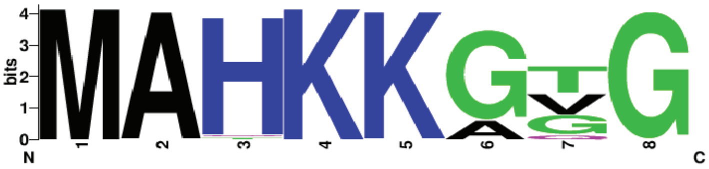


**Fig. S3. Sequence logo of bL27 proteins generated from the 100 most diverse PRK05435 sequences.** The stack and letter heights indicates positional information content and amino-acid frequency.

**Table S1. Strains and plasmids used for this work**

| **Strain or plasmid** | **Description** | **Source or reference** |
|---|---|---|
| MG1655 $\Delta$*tolC*::cat | *tolC* deletion<br>Referred to as $\Delta$*tolC* | (*34*) |
| IW312 $\Delta$*rpmA*::Kan | bL27 deletion strain | (*24*) |
| X90 | *E. coli* strain used for *ssrA* modification in the genome | Gift from C. Hayes |
| X90 *ssrA*6HisD Amp$^{R}$ | *ssrA* in genome encodes 6HisD with Amp$^{R}$ | Gift from C. Hayes |
| X90 *ssrA*6HisD Amp$^{R}$ $\Delta$*rpmA*::kan | *ssrA* in genome encodes 6HisD with Amp$^{R}$ with *rpmA* deletion | This work |
| MG1655 $\Delta$*ssrA*::cat | *ssrA* deletion strain<br>Referred to as $\Delta$*ssrA* | (*34*) |
| MG1655 $\Delta$*tolC*::cat $\Delta$*rpmA*::kan | *tolC, rpmA* deletion<br>Host for pL27 plasmids | This work |
| MG1655 $\Delta$*tolC* $\Delta$*ssrA*::cat $\Delta$*rpmA*::kan | *tolC*, tmRNA, and bL27 deletion strain | This work |

| | | |
|---|---|---|
| MG1655 Δ*tolC*::cat Δ*rpmA*::kan pL27 AHKKAGG $Amp^R$ | Wild-type bL27 is expressed via pL27 in bL27 deletion strain<br>Referred to as Δ*rpmA* pL27 | This work |
| MG1655 Δ*tolC*::cat Δ*rpmA*::kan pL27 AaKKAGG $Amp^R$ | H3A in N-terminal bL27 sequence<br>Referred to as AaKKAGG | This work |
| MG1655 Δ*tolC*::cat Δ*rpmA*::kan pL27 AHaKAGG $Amp^R$ | K4A in N-terminal bL27 Sequence<br>Referred to as AHaKAGG | This work |
| MG1655 Δ*tolC*::cat Δ*rpmA*::kan pL27 AHKaAGG $Amp^R$ | K5A in N-terminal bL27 sequence<br>Referred to as AHKaAGG | This work |
| MG1655 Δ*tolC*::cat Δ*rpmA*::kan pL27 AHKKAaG $Amp^R$ | G7A in N-terminal bL27 sequence<br>Referred to as AHKKAaG | This work |
| MG1655 Δ*tolC*::cat Δ*rpmA*::kan pL27 AHKKAGa $Amp^R$ | G8A in N-terminal bL27 sequence<br>Referred to as AHKKAGa | This work |
| MG1655 Δ*tolC*::cat Δ*rpmA*::kan pL27 AHKKAaa $Amp^R$ | G7A G8A in N-terminal bL27 sequence<br>Referred to as AHKKAaa | This work |
| MG1655 Δ*tolC*::cat Δ*rpmA*::kan pL27 MHKKAGG $Amp^R$ | ΔA2 in N-terminal bL27 sequence<br>Referred to as MHKKAGG | This work |
| MG1655 Δ*tolC*::cat Δ*rpmA*::kan pL27 aKKAGG $Amp^R$ | ΔA2 H3A in N-terminal bL27 sequence<br>Referred to as aKKAGG | This work |
| MG1655 Δ*tolC*::cat Δ*rpmA*::kan pL27 AaaaAGG $Amp^R$ | H3A K4A K5A in N-terminal bL27 sequence<br>Referred to as AaaaAGG | This work |
| MG1655 Δ*tolC*::cat Δ*rpmA*::kan pL27 MKAGG $Amp^R$ | -3 truncation in N-terminal bL27 sequence<br>Referred to as MKAGG | This work |
| MG1655 Δ*tolC*::cat Δ*rpmA*::kan pL27 G $Amp^R$ | -6 truncation in N-terminal bL27 sequence<br>Referred to as G | This work |
| MG1655 Δ*tolC*::cat Δ*rpmA*::kan vector only $Amp^R$ | bL27 is deleted in the genome and contains the empty plasmid<br>(No *rpmA* gene in the plasmid)<br>Referred to as Δ*rpmA* | This work |

| | | |
|---|---|---|
| MG1655 Δ*tolC* Δ*ssrA*::cat Δ*rpmA*::kan pL27 AHKKAGG $Amp^R$ | *ssrA* deletion in strain with pL27<br>Referred to as Δ*rpmA* Δ*ssrA* pL27 | This work |
| MG1655 Δ*tolC* Δ*ssrA*::cat Δ*rpmA*::kan pL27 AaaaAGG $Amp^R$ | *ssrA* deletion in strain with pL27 AaaaAGG<br><br>Referred to as Δ*rpmA* Δ*ssrA* pL27 AaaaAGG | This work |
| MG1655 Δ*tolC* Δ*ssrA*::cat Δ*rpmA*::kan vector only $Amp^R$ | *ssrA* deletion in Δ*rpmA* background, contains empty plasmid<br>Referred to as Δ*rpmA* Δ*ssrA* | This work |
| pL27 AHKKAGG $Amp^R$ | Plasmid expressing Wild-type bL27 N-terminal sequence | (*25*) |
| pL27 MKAGG $Amp^R$ | Plasmid expressing -3 truncation in bL27 N-terminal sequence | (*25*) |
| pL27 G $Amp^R$ | Plasmid expressing -6 truncation in bL27 N-terminal sequence | (*25*) |
| Vector only $Amp^R$ | No *rpmA* gene in the pL27 plasmid | (*25*) |
| pCP20 | Used for Flp catalyzed excision of CAT cassette | (*56*) |
| JW2667 | Contains plasmid pCA24N expressing *E.coli alaS* | (*57*) |
| JW1865 | Contains plasmid pCA24N expressing *E.coli argS* | (*57*) |
| JW1855 | Contains plasmid pCA24N expressing *E.coli aspS* | (*57*) |
| JW0913 | Contains plasmid pCA24N expressing *E.coli asnS* | (*57*) |
| JW0515 | Contains plasmid pCA24N expressing *E.coli cysS* | (*57*) |

| | | |
|---|---|---|
| JW0666 | Contains plasmid pCA24N expressing *E.coli glnS* | (*57*) |
| JW2395 | Contains plasmid pCA24N expressing *E.coli gltX* | (*57*) |
| JW3531 | Contains plasmid pCA24N expressing *E.coli glyQ* | (*57*) |
| JW3530 | Contains plasmid pCA24N expressing *E.coli glyS* | (*57*) |
| JW2498 | Contains plasmid pCA24N expressing *E.coli hisS* | (*57*) |
| JW0024 | Contains plasmid pCA24N expressing *E.coli ileS* | (*57*) |
| JW0637 | Contains plasmid pCA24N expressing *E.coli leuS* | (*57*) |
| JW4090 | Contains plasmid pCA24N expressing *E.coli lysU* | (*57*) |
| JW2101 | Contains plasmid pCA24N expressing *E.coli metG* | (*57*) |
| JW5277 | Contains plasmid pCA24N expressing *E.coli* pheS | (*57*) |
| JW1703 | Contains plasmid pCA24N expressing *E.coli* pheT | (*57*) |
| JW0190 | Contains plasmid pCA24N expressing *E.coli proS* | (*57*) |
| JW0876 | Contains plasmid pCA24N expressing *E.coli serS* | (*57*) |

| | | |
|---|---|---|
| JW1709 | Contains plasmid pCA24N expressing *E.coli thrS* | (*57*) |
| JW3347 | Contains plasmid pCA24N expressing *E.coli trpS* | (*57*) |
| JW1629 | Contains plasmid pCA24N expressing *E.coli tyrS* | (*57*) |
| JW4215 | Contains plasmid pCA24N expressing *E.coli* v*alS* | (*57*) |
| JW2502 | Contains plasmid pCA24N expressing *E.coli* n*dk* | (*57*) |
| JW3249 | Contains plasmid pCA24N expressing *E.coli fmt* | (*57*) |
| JW0867 | Contains plasmid pCA24N expressing *E.coli infA* | (*57*) |
| JW3137 | Contains plasmid pCA24N expressing *E.coli infB* | (*57*) |
| JW5829 | Contains plasmid pCA24N expressing *E.coli infC* | (*57*) |
| JW3302 | Contains plasmid pCA24N expressing *E.coli fusA* | (*57*) |
| JW3301 | Contains plasmid pCA24N expressing *E.coli tufA* | (*57*) |
| JW0165 | Contains plasmid pCA24N expressing *E.coli tsf* | (*57*) |
| JW1202 | Contains plasmid pCA24N expressing *E.coli prfA* | (*57*) |

| | | |
|---|---|---|
| JW5847 | Contains plasmid pCA24N expressing *E.coli prfB* | (*57*) |
| JW5873 | Contains plasmid pCA24N expressing *E.coli prfC* | (*57*) |
| JW0167 | Contains plasmid pCA24N expressing *E.coli frr* | (*57*) |
| pDHFR | Expresses DHFR off a T7 promoter; $Amp^R$ | NEB |
| *E. coli* BL21 (DE3) pET28 *smpB* | expresses his-tagged SmpB | (*34*) |
| *E. coli* BL21 (DE3) pBP-T7RNAP $Amp^R$ | expresses his-tagged T7 RNA polymerase | Gift from Bevilacqua lab |

**Table S2. Primers used for this work**

| Oligonucleotide | SEQUENCE | Description | Reference or source |
|---|---|---|---|
| F1 | GGCTGGCGGCTCCACACGTA | Vector primer | This work |
| R1 | CATTTGAAATCTCTCCTCAGGTC TTAAGA | Vector primer | This work |
| F2 | ACGTAACGGTCGCGATTCAG | Vector primer | This work |
| R2 | GCCTTTTTATGTGCCATTTGAAA TCTCT | Vector primer | This work |
| AaKKAGG | TCTTAAGACCTGAGGAGAGATT TCAAATGGCAgctAAAAAGGCTG GCGGCTCCACACGTA | Assembled with F1R1 amplified vector | This work |
| AHaKAGG | TCTTAAGACCTGAGGAGAGATT TCAAATGGCACATgcaAAGGCTG GCGGCTCCACACGTA | Assembled with F1R1 amplified vector | This work |
| AHKaAGG | TCTTAAGACCTGAGGAGAGATT TCAAATGGCACATAAAgcgGCTG GCGGCTCCACACGTA | Assembled with F1R1 amplified vector | This work |
| AHKKAaG | AGAGATTTCAAATGGCACATAA AAAGGCTgccGGCTCCACACGTA ACGGTCGCGATTCAG | Assembled with F2R2 amplified vector | This work |
| AHKKAGa | AGAGATTTCAAATGGCACATAA AAAGGCTGGCgccTCCACACGTA ACGGTCGCGATTCAG | Assembled with F2R2 amplified vector | This work |

| | | | |
|---|---|---|---|
| AHKKAaa | AGAGATTTCAAATGGCACATAAAAAGGCTgccgccTCCACACGTAACGGTCGCGATTCAG | Assembled with F2R2 amplified vector | This work |
| MHKKAGG | TCTTAAGACCTGAGGAGAGATTTCAAATGCATAAAAAGGCTGGCGGCTCCACACGTAACG | Assembled with F1R1 amplified vector | This work |
| aKKAGG | TCTTAAGACCTGAGGAGAGATTTCAAATGgctAAAAAGGCTGGCGGCTCCACACGTAACG | Assembled with F1R1 amplified vector | This work |
| aKAGG | TCTTAAGACCTGAGGAGAGATTTCAAATGgcaAAGGCTGGCGGCTCCACACGTAACGGTC | Assembled with F1R1 amplified vector | This work |
| AaaaAGG | TCTTAAGACCTGAGGAGAGATTTCAAATGGCAgctgcagcgGCTGGCGGCTCCACACGTA | Assembled with F1R1 amplified vector | This work |
| metZ_for | TAATACGACTCACTATAGGGCGCGGGGTGGAGCAGC | Amplify *metZ* from genomic DNA with T7 promoter | This work |
| metZ_rev | TGGTTGCGGGGGCCGGATTTGAAC | Amplify *metZ* from genomic DNA | This work |
| *α-ssrA* | TTAAGCTGCTAAAGCGTAGTTTTCGTCGTTTGCGACTA | Primer used to block background tagging from *ssrA* | (*34*) |
